# Finding domain-general metacognitive mechanisms requires using appropriate tasks

**DOI:** 10.1101/211805

**Authors:** Eugene Ruby, Nathan Giles, Hakwan Lau

**Author notes:** **Correspondence should be addressed to:** Eugene Ruby UCLA Psychology Department 1285 Franz Hall, Box 951563 Los Angeles, CA 90095-1563.

## Abstract

An important yet unresolved question is whether or not metacognition consists of domain-general or domain-specific mechanisms. While most studies on this topic suggest a dissociation between metacognitive abilities at the neural level, there are conflicting reports at the behavioral level. Specifically, while McCurdy et al. (2013) found a positive correlation between metacognitive efficiency for visual perception and memory, Baird et al. (2013) didn’t find this correlation. One possible explanation for this disparity is that the former included two-alternative-forced choice (2AFC) judgments in both their visual and memory tasks, whereas the latter used 2AFC for one task and yes/no (YN) judgments for the other. In support of this hypothesis, we ran two online experiments meant to mirror McCurdy et al. (2013) and Baird et al. (2013) with considerable statistical power (n=100), and replicated the main findings of both studies. This suggests the finding of McCurdy et al (2013) was not a ‘fluke’ (i.e. false positive). In a third experiment with the same sample size, which included YN judgments for both tasks, we did not find a correlation between metacognitive functions, suggesting that the conflict between McCurdy et al. (2013) and Baird et al. (2013) stemmed from the use of YN judgments in the latter study. Our results underscore the need to avoid conflating 2AFC and YN judgments, which is a common problem.

## Introduction

Metacognition is the crucial cognitive ability that enables us to monitor and regulate our own mental processes. In experiments, one way to quantify metacognition is to assess the trial-by-trial correspondence between confidence and accuracy in behavioral tasks. An important question that remains unclear is whether different types of metacognition, like metacognition for visual perception and memory, depend on distinct, domain-specific neurocognitive mechanisms, or alternatively on a single, domain-general system that supports metacognition for all mental faculties.

Interestingly, while empirical studies on this issue mostly cohere in suggesting a dissociation between metacognitive functions at the neural level (Baird et al., 2013; McCurdy et al., 2013; Fleming et al., 2014; Baird et al., 2015; Morales et al., 2017), findings are somewhat conflicted at the behavioral level. Specifically, while McCurdy et al. (2013) reported a positive correlation between metacognitive efficiency for memory (i.e., metamemory) and visual perception, Baird et al. (2013) found no correlation.

One possible explanation is a difference in statistical power between the studies, or perhaps a lack of power in both. Therefore we attempted to replicate both experiments, but we ran them online to acquire larger sample sizes for increased power.

Another issue is that both McCurdy et al. (2013) and Baird et al. (2013) didn’t use the same stimulus type across the two tasks (both studies used circles with gratings for the visual task, but words for the memory task). In general, this may have made it harder to detect an effect. To address this potential confound, we matched the stimulus type across tasks in our experiments.

While the above issues may have contributed to the conflicting results, we hypothesize that the most important factor was a subtle (yet crucial) difference between the two studies, one that Baird et al. (2013) themselves suggest may have contributed. While McCurdy et al. (2013) required two-alternative forced choice (2AFC) discrimination judgments for both tasks, Baird et al. (2013) required 2AFC judgments for the visual task but yes/no (YN) judgments for the memory task. In 2AFC task, in each trial participants are presented with a pair of stimuli, and they are required to identify the spatial or temporal arrangements of the pair, e.g. old word on the left, new word on the right, or vice versa (as was done in McCurdy et al 2013). In YN judgments, subjects answer a binary question about a single stimulus in each trial, e.g. whether a word is new or old (as was done in Baird et al 2013). Because 2AFC and YN tasks presumably involve rather different mechanisms, we suspect that confusing and mixing them together in the same experiment is an important and yet often ignored confound. To tackle this issue, we assessed in a series of experiments as to whether adopting the same 2AFC judgments for both tasks as in McCurdy et al. (2013) is crucial for assessing correlation of metacognitive efficiency across domains (memory vs. vision).

## Methods (for all experiments)

### Subjects

In each of the following experiments, one hundred healthy subjects were recruited using Amazon Mechanical Turk’s task hosting service and the experiment was conducted through the Internet (using the same service). Eligibility was determined by the subjects and listed in an online advertisement and consent form for the study; in order to take part in the experiment, all subjects were required to have normal or corrected-to-normal vision (e.g., glasses or contacts) and no history of any psychiatric or neurological illnesses or seizures. All subjects provided consent to participate. Subjects were compensated $4 for completion of the study, with the possibility of receiving a $1 bonus if their performance was higher than that of the previous participant.

### Behavioral Tasks and Stimuli

As in McCurdy et al. (2013) and Baird et al. (2013), all subjects completed both a visual task and a memory task.

In all of our experiments, subjects completed 120 trials of the visual task. The task progression differed depending on whether 2AFC or YN judgments were included. On each trial of the visual 2AFC task (used in Experiments 1 and 2), participants were presented with two different clusters of circles of varying sizes for 475 milliseconds, followed by a blank screen for 500 milliseconds. Next, participants were given 2.25 seconds to indicate whether the average size of the circles was larger in the left or right cluster from the previous screen (by pressing 1 for left or 2 for right). Another screen followed giving subjects 2.25 seconds to indicate how confident they were in their discrimination judgment by pressing a number from 1 to 6, with 6 being extremely confident and one being not at all confident. Finally, a second blank screen was displayed for 1.5 seconds, before the next trial begins. This entire process constituted one trial of the visual 2AFC task (Figure 1a). As for the visual YN task (used in Experiment 3), subjects were shown 10 examples of clusters of circles of “medium” size before the start of the task. During the task, subjects were shown one cluster of circles and subsequently asked to indicate whether this cluster contained circles that were on average bigger or smaller than those of the medium cluster. One trial on the visual YN task was in every other way identical to one trial on the visual 2AFC task (Figure 1b).

**Figure 1:**
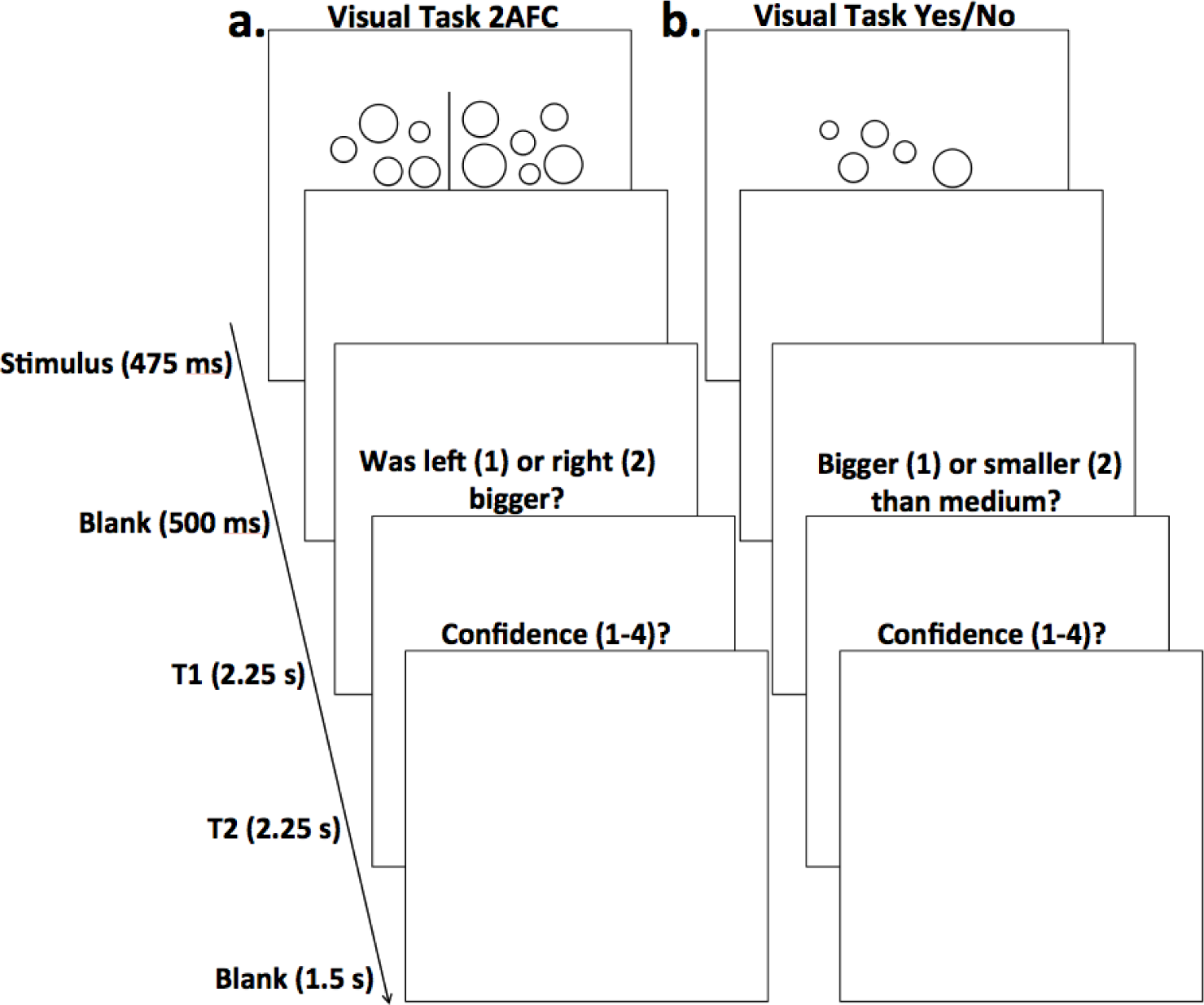
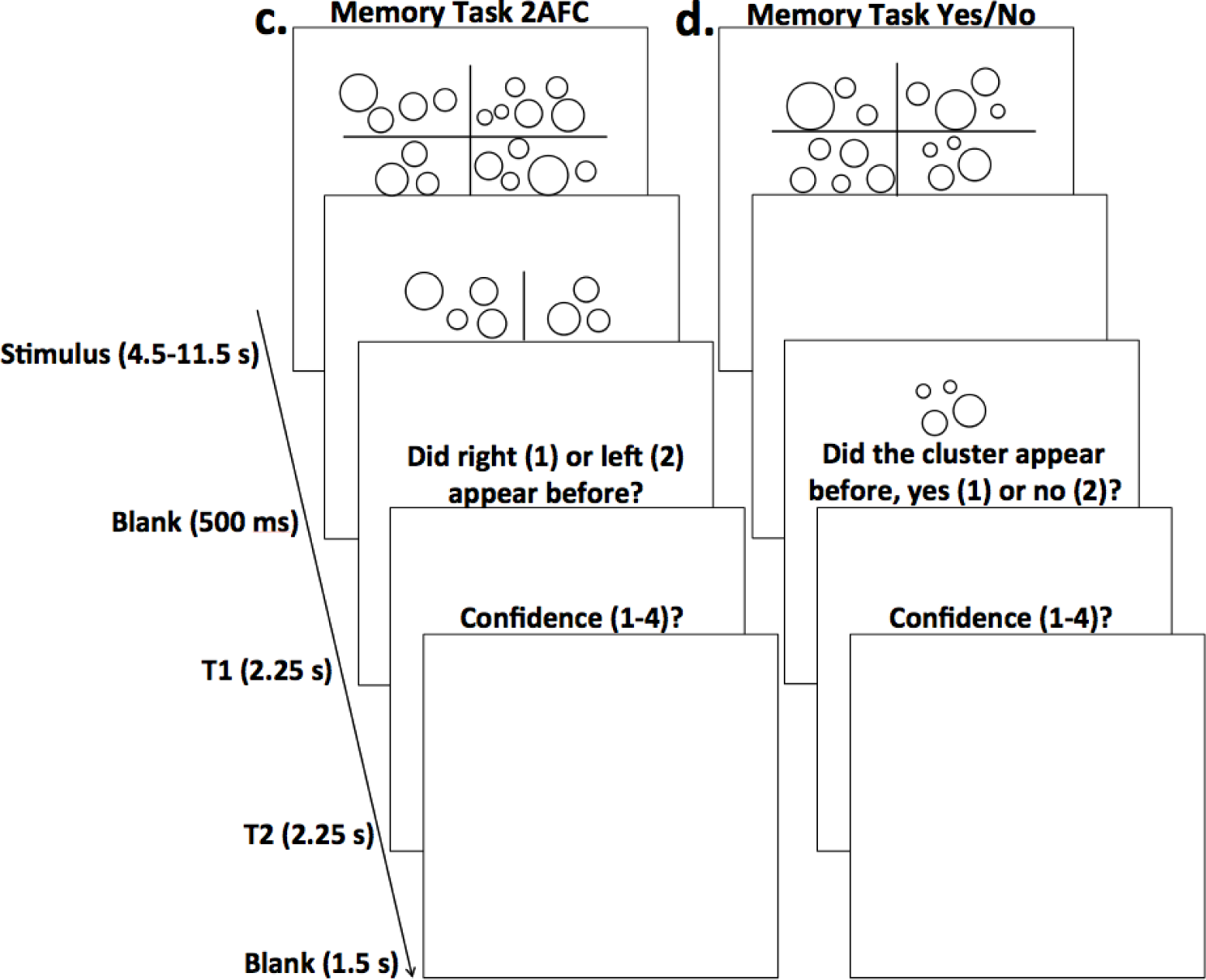
Timeline for individual trials on the various tasks. **(a) Visual task with 2AFC judgments.** Subjects were shown two clusters of circles, followed by a blank screen, and were then prompted to discriminate whether the cluster with (on average) larger circles was on the left or right (T1). They then rated how confident they were on a scale of 1-6 (with 1 being not at all confident and 6 being completely confident) in their discrimination judgment (T2). A blank screen ended the trial. **(b) Visual task with YN judgments.** Each visual task trial requiring YN judgments was identical to trials for the 2AFC visual task, except that subjects were shown only one cluster, and were later prompted to discriminate whether the cluster had (on average) larger circles than a cluster of “medium” size (T1). **(c) Memory task with 2AFC judgments.** Subjects were shown four clusters of circles, followed by a blank screen. Next, two clusters were presented, one from the previous stimulus presentation and one new cluster; participants were instructed to pick the cluster that appeared before (T1). As with the visual tasks, they then gave confidence judgments on a scale of 1-6 (T2). A blank screen ended the trial. **(d) Memory task with YN judgements.** Each memory task trial requiring YN judgments was identical to trials for the 2AFC memory task, except that only one cluster was presented during the discrimination portion of the task and subjects were instructed to determine whether it had appeared on the previous stimulus presentation or not (T1).

As in McCurdy et al. (2013) and Baird et al. (2013), we used staircase procedures for our visual tasks in order to keep discrimination performance relatively constant. We aimed to keep subjects’ basic task performance at around a 75% threshold, where psychophysical measurements are most efficient. For the visual 2AFC task, the difference in average size of each cluster of circles (and thus, also the sizes of the circles within each cluster) was generated on each trial by randomly selecting from one of two staircase procedures. The first was a two-up-one-down staircase; any one incorrect response to a discrimination judgment (on any given trial) caused the difference in average size of each cluster of circles to increase by 1 pixels, and any two consecutive correct responses resulted in this difference in average size to decrease by 1 pixel. The second staircase used a three-up-one-down procedure; any one incorrect response to a discrimination judgment caused the difference in average size of each cluster of circles to increase by 1 pixels, and any three consecutive correct responses (on back to back trials) resulted in this difference in average size to decrease by 1 pixel. The baseline difference in average size of each cluster of circles in terms of radii was 6 pixels, the minimum difference allowable was 1 pixel, and the maximum was 11 pixels. For the visual YN task, we randomly selected from one of two one up one down staircases on every trial; for each staircase, any correct response caused the difference between a trial cluster’s average radius and the “medium” cluster’s average radius to decrease by 1 pixel, and any incorrect response led to this difference increasing by 1 pixel. The baseline difference was 4 pixels, the minimum difference allowable was 1 pixel, and the maximum was 7 pixels. For both visual tasks, the positions of the circles within each cluster were not allowed to overlap, and were kept at a set minimum distance from any other circles in that cluster. Similarly, they were also not allowed any further than a set maximum distance from at least one neighboring circle. Given these restrictions, the positions of the circles within each cluster were generated in a pseudo-random manner.

As with the visual tasks, for each of our memory tasks subjects completed 120 trials. The task progression differed depending on whether 2AFC or YN judgments were included. On each trial of the memory 2AFC task (used in Experiment 1), subjects were first shown a stimulus presentation of four clusters of circles of varying sizes to remember for anywhere from two to eight seconds, followed by a blank screen for 500 milliseconds. Next, subjects were shown two circle clusters, one from the previous stimulus presentation and one new cluster, and had 2.25 seconds indicate which of the two was previously presented (by pressing 1 for the left cluster or 2 for the cluster on the right). Subjects were then given 2.25 seconds to indicate how confident they were in their discrimination judgment by pressing a number from 1 to 6, with 6 being extremely confident and 1 being not at all confident (i.e., metamemory confidence judgments). A blank screen followed for 1.5 seconds to complete a single trial. The following three trials each contained the previous three steps (discrimination judgment, confidence judgment, and post-trial interval), but didn’t include a new stimulus presentation. All four trials just described taken together constituted one mini-block for the memory 2AFC task (Figure 1c). The memory YN task (used in Experiments 2 and 3) was identical to the memory 2AFC task in every way, except that subjects were shown a single cluster of circles (instead of two) during the discrimination portion of the task and asked to indicate whether or not they remembered seeing the cluster on the study list at the beginning of a given trial with a “yes” or “no” response (Figure 1d).

As with the visual tasks, we used staircase procedures to keep discrimination performance at around 75%. Both the memory 2AFC and YN tasks used the same staircase procedure. For the presentation of the study list at the beginning of each mini-block, the duration for which the stimuli were shown was determined using a variation of a four-up-two-down staircase procedure; for every mini-block with perfect discrimination performance (i.e., all four discrimination judgments in a given mini-block were answered correctly), the time that the study list was shown on the following mini-block decreased by 500 milliseconds. On the other hand, for every mini-block in which the participant responded incorrectly to at least two (of the four) trials, the time that the study list was shown on the following mini-block increased by 500 milliseconds. Otherwise, the time the study list was shown on the next trial remained the same. The baseline study list duration at the beginning of the experiment was eight seconds. The minimum and maximum allowable study list durations were 4.5 and 11.5 seconds, respectively.

For each visual or memory task, 5 percent of all trials were catch trials, which helped to make sure that participants were actively attending to the task, and furthermore that they correctly understood the task instructions. Each catch trial consisted of a single trial with a very easy discrimination judgment. If subjects performed poorly on any two catch trials, they were told that they hadn’t performed well on some of the easier trials and were asked whether or not they want to continue the experiment. Moreover, subjects that didn’t give responses in time for four consecutive trials were given a warning, indicating that they should pay closer attention to the task, and that they would not be able to continue the experiment if they failed to do so. Following this, if subjects again didn’t respond on four consecutive trials, their participation in the experiment was terminated immediately. The order of the tasks was counterbalanced across subjects. The entire experiment took approximately one hour to complete. For every study participant, all of the tasks were completed in one session. As remaining stationary and on task for one hour might induce some boredom or minor physical discomfort, subjects were given two short breaks throughout each of the two tasks in order to ensure they retained a sufficient level of comfort and focus.

All stimuli and behavioral tasks were created using JavaScript by utilizing jsPsych, which is a JavaScript library used to design and run psychological experiments. The stimuli in the tasks were scaled off of the size of each subject’s computer screen, which ensured that the size and position of all stimuli would be the same for all subjects, despite the fact that computer screen size probably varied from participant to participant.

### Data Analysis

To assess metacognitive sensitivity for each modality, we utilized the bias-free psychophysical measure, meta-d’ (Maniscalco and Lau, 2012), which measures how well participants can differentiate between correct and incorrect judgments on a trial-by-trial basis (Fleming & Lau, 2014). We then divided subjects’ meta-d’ by their d’ score (basic discrimination performance) to obtain meta-d’/d’ (M Ratio), a measure of metacognitive efficiency (i.e., a participant’s metacognitive sensitivity given a specific level of basic task performance). M Ratio, which helps to control for the effects of basic task performance (McCurdy et al. 2013), was also the primary measure of interest in both McCurdy et al. (2013) and Baird et al. (2013); i.e., correlational analyses in these studies assessed the relationship between M Ratio for visual metacognition and metamemory. After data collection, we similarly performed within-subjects correlation analyses to assess relationships between visual metacognition and metamemory.

For each of our experiments below we focus on the results of the correlation analyses after removing influential outliers. To identify such outliers Cook’s D (Cook, 1977) was calculated for all data points; if Cook’s D for any subject exceeded the standard threshold recommendation of 4/(n-k-1),where n is the sample size and k is the number of independent variables, then that subject’s data was labeled an outlier with an influential effect on the regression line, and it was removed from the analysis. Notably, however, in all experiments we found similar patterns of statistical significance and results, both regardless whether outliers were removed or not (unless otherwise noted in specific cases below).

## Results

### Experiment 1

As in McCurdy et al. (2013), both tasks in our Experiment 1 included 2AFC judgments (see Methods). Ninety-six subjects were included in analyses after subjects exclusion, giving our experiment more statistical power than McCurdy et al. (2013), who included only 34 subjects. We found a significant positive correlation between M Ratio for visual and memory metacognition (r = 0.3067, p = 0.0024; Figure 2). Of note, this significant correlation was present also without removing influential outliers, with a numerically weaker strength (r = 0.2470, p = .0132).

**Figure 2:**
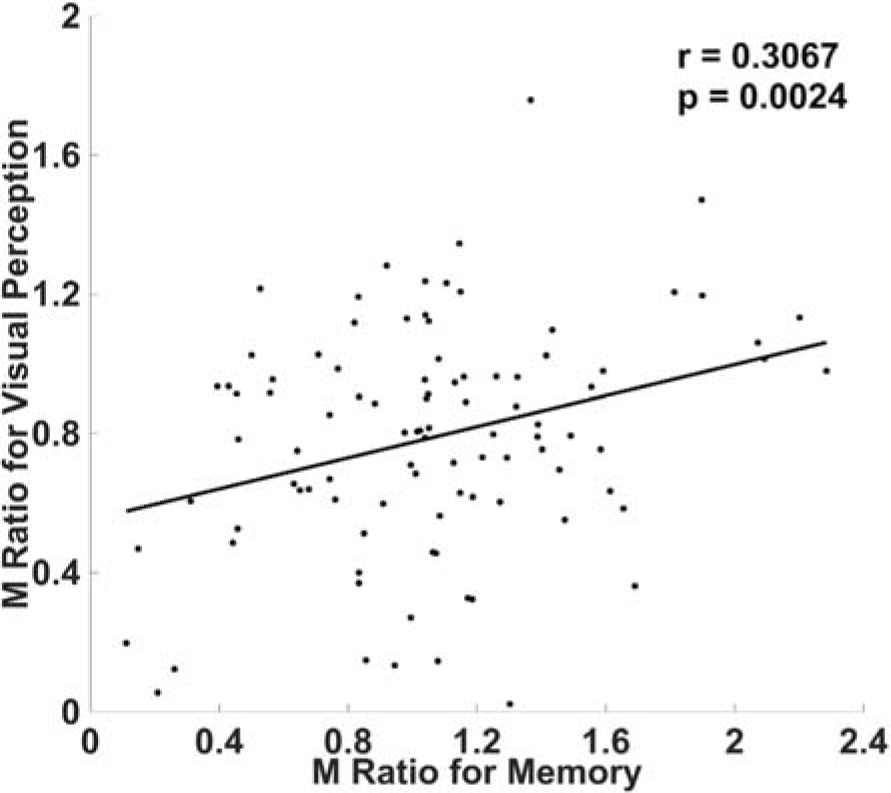
Correlation between visual and memory metacognitive efficiency when when both tasks involved 2AFC. As in McCurdy et al. (2013), we found a significant positive correlation across subjects, between visual and memory metacognitive efficiency when both tasks involved 2AFC rather than YN judgments. Metacognitive efficiency was quantified using M Ratio, a detection theoretic measure of metacognitive efficiency that accounts for fluctuations in task performance (see Methods).

### Experiment 2

Given our replication of the main finding of McCurdy et al. (2013) in Experiment 1, in our Experiment 2 we changed the discrimination judgments for the memory task from 2AFC to YN judgments while leaving everything else from our first experiment unchanged (see Methods), and attempted to mimic the set up by Baird et al. (2013). The idea was to see if using this asymmetric design (with one task being 2AFC and the other being YN), one could still observe a significant correlation between metacognitive efficiencies when we have enough subjects.

Ninety-three subjects were included in analyses after subjects exclusion, giving this study more statistical power than Baird et al. (2013), who included 52 subjects. As in Baird et al. (2013), we failed to find a correlation between M Ratio for memory and visual metacognition (r = 0.0739, p = 0.4815; Figure 3). In short, we replicated the behavioral result from Baird et al. (2013).

**Figure 3:**
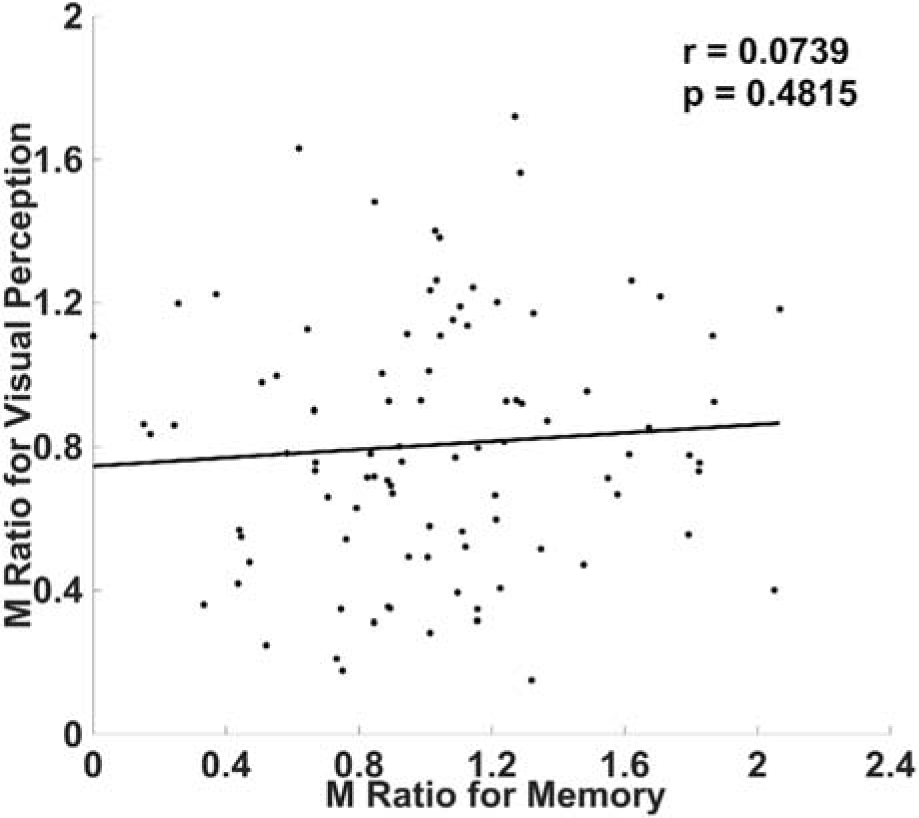
No correlation in metacognitive efficiency when the visual task involved 2AFC judgments and the memory task involved YN judgments. As in Baird et al. (2013), we didn’t find a correlation between visual and memory metacognitive efficiency when the visual task involved 2AFC judgments and the memory task involved YN judgments. As in Experiment 1, metacognitive efficiency was quantified using M Ratio (see Methods).

#### Across Experiments Analyses

To further probe whether the difference in behavioral results between the two studies can be consistently replicated, we converted Pearson correlation coefficient values for each of the two experiments to z values (as z values, unlike r values, are normally distributed and can therefore be compared) using Fisher’s R to Z transformation (Fisher, 1915). Z tests were then run (Cohen & Cohen, 1983) to test the hypothesis (one sided) that the correlation between M Ratio for visual and memory metacognition decreased significantly from our Experiment 1 to Experiment 2; this hypothesis was supported (z = 1.77, p = .0394). Thus, not only did we find different results when using distinct types of discrimination judgments across the two experiments, this difference was also significant under a direct comparison.

### Experiment 3

The results of our first two experiments raise the question of what exactly it is about the distinction in judgment type that causes differing results. One possibility is that observing a correlation depends on using the same judgment type (2AFC vs YN) across tasks. For example, both our Experiment 1 and McCurdy et al. (2013) found significant correlations when using the same type of discrimination judgment for both tasks (2AFC judgments); if the correlations depend on using the same type of judgment across tasks, perhaps we should also find that utilizing YN judgments for both tasks will similarly reveal a significant correlation.

Alternatively, our Experiment 2 and Baird et al. (2013) may have failed to find a correlation because YN judgments are somewhat more complicated than 2AFC judgments. 2AFC judgments involve two stimulus alternatives being directly compared based on their ‘familiarity’ (or some other feature reflecting a conscious memory trace), whereas YN judgments involve only one stimulus being present and no other stimulus directly accessible to compare with the present stimulus. As such, performing a YN judgement requires a stable criterion as to what count as ‘familiar’. This may place a higher demand on working memory given that the participant likely has to compare the present stimulus to other stimuli that have recently been encountered but are no longer present.

With these possibilities in mind, we ran a third experiment, using YN judgments for not only the memory task (as in Experiment 2), but also for the visual task (see Methods). Ninety-four subjects were included in analyses after subjects exclusion We did not find a correlation between M Ratio for metamemory and visual metacognition (r = 0.0934, p = 0.3707; Figure 4). Taken alongside the results of our first two experiments, this suggests the conflicting behavioral findings from Rounis et al. (2013) and Bor et al. (2013) reflects the fact that YN judgments are the culprit.

**Figure 4:**
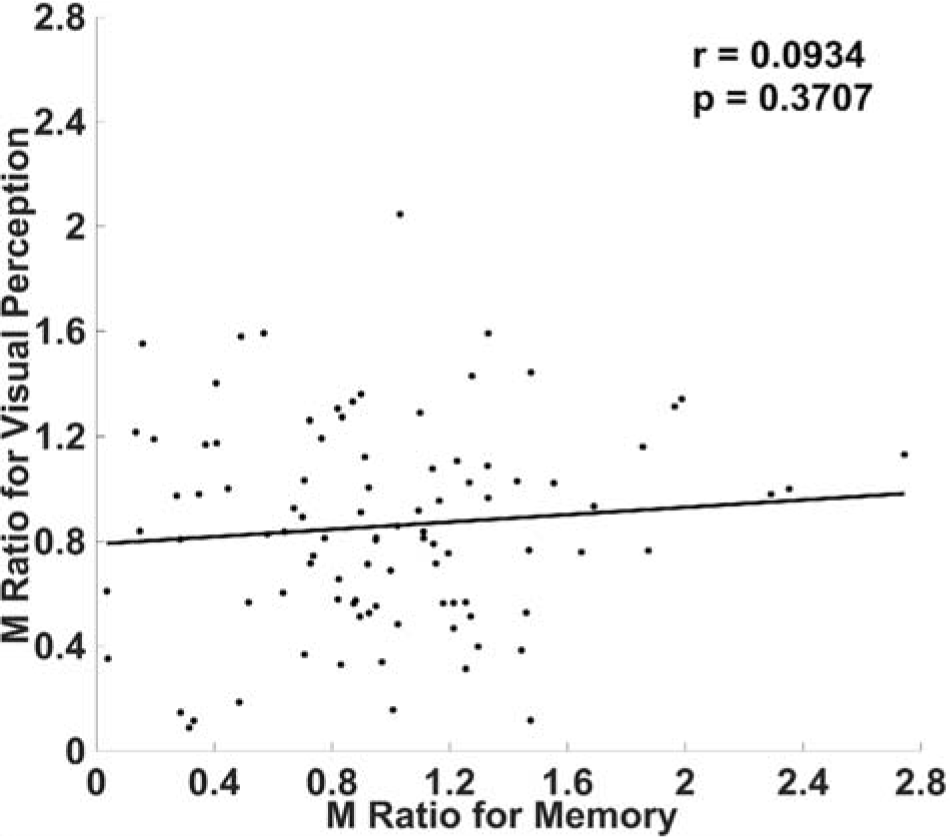
No correlation between visual and memory metacognitive efficiency when both tasks involved YN judgments. We did not find a correlation between visual and memory metacognitive efficiency when both tasks involved YN rather than 2AFC judgments. Metacognitive efficiency was quantified using M Ratio (see Methods).

#### Further Analyses

If YN judgments are in fact somehow limiting our ability to reveal metacognitive correlations than 2AFC judgments, what exactly is it about the former that makes it so? Above we suggested the possibility that YN judgments might be more demanding in terms of criterion maintenance, and possibly other higher cognitive processes. Further dissecting this proposal, we might ask if this added complexity is true for both “yes” and “no” responses. Of relevance, several previous studies suggest that metacognitive sensitivity is lower for “no” responses than for “yes” responses (Kanai et al., 2010; Maniscalco & Lau, 2011).

We ran further analyses, assessing meta-d’ separately for trials where the participants made a ‘yes’ or a ‘no’ response. Based on the data from Experiment 3, we confirmed that meta d’ for “yes” responses significantly correlated across tasks (r = .2364, p = .0179), but this wasn’t the case for meta d’ for “no” responses (r = .1453, p = .1493). Therefore, it appears that the lack of correlations observed in our Experiment 3, as well as in our Experiment 2 and Baird et al. (2013), were due specifically to some idiosyncratic nature of metacognition after “no” responses.

In general, it is known that metacognition after “no” responses may be less efficient (Kanai et al., 2010; Maniscalco & Lau, 2011). This was numerically true for both our Experiment 2 memory task, for which meta d’ for “yes” responses was significantly greater than for “no” responses (the means were

1.511 and 0.990 respectively, t(99) = −4.475, p > .001), and for our Experiment 3 memory task (the means were 1.295 and 1.106 for meta-d’ after ‘yes’ and ‘no’ responses respectively), although the latter didn’t reach statistical significance, t(99) = −1.255, p = .213. Conversely for our Experiment 3 visual task, meta d’ for “yes” responses was lower than for “no” responses (the means were 1.285 and 1.169 respectively; the difference wasn’t significant, t(99) = .594, p = .554). Based on these results, we further tested the possibility that meta-d’ for “no” responses is problematic for assessing cross-domain metacognitive correlations, only when it is lower than meta-d’ for “yes” responses. In support of this possibility, we found that in Experiment 3, meta d’ for “yes” responses on the memory task was significantly associated with (overall) meta d’ for the visual task in our Experiment 3 (r = .2131, p = .0333), meaning that including the metacognitive measure after “no” responses was not a problem for revealing metacognitive correlations. Likewise, in Experiment 2 we also tested for an association between meta d’ for “yes” responses on the memory task and (overall) meta d’ for the visual task and found a strong trend towards significance (r = .1890, p = .0596). Further supporting the above suggestion, we found that meta d’ for “yes” responses on the visual task wasn’t significantly related to (overall) meta d’ for the memory task in our Experiment 3 (r = .1640, p = .1031), where meta-d’ for “no” responses wasn’t smaller than meta-d’ for “yes” responses. In sum, it seems that including meta-d’ for “no” responses is only limiting our ability to reveal across-domain metacognition when it is lower than meta-d’ for “yes” responses. Below we discuss what this may mean, in terms of mechanisms.

## Discussion

We showed that requiring 2AFC judgments for both a visual task and memory task allowed us to show a positive correlation between visual and memory metacognition (Experiment 1), whereas this correlation significantly decreased and ultimately disappeared when requiring 2AFC for one task and YN judgments for the other (Experiment 2). We also found no correlation when requiring YN judgments for both tasks (Experiment 3). Together, these results suggest that our failure to find a correlation in Experiments 2 and 3 resulted specifically from YN judgments introducing more ambiguity into the decision process than 2AFC. Further analyses indicated that specifically, it is metacognition after “no” responses that may be the problem, especially in the memory task.

What is it specifically about “no” responses in the memory task that might have caused the observed results? One subtle difference between our YN memory and visual tasks is that the former is what has been called a ‘true’ detection task (the goal of which is to detect a stimulus presence from a stimulus absence) whereas the latter is a ‘pseudo’ detection task (in which one detects the presence of a stimulus *feature* while the absence case contains a similar level of stimulus energy despite the lack of that particular feature). This distinction was introduced by Maniscalco and Lau (2011), who reported that meta d’ for “no” responses was lower than for “yes” responses on task in which the target-present condition involved the presence of some physical feature (solid dots), which carried additional stimulus energy.

However, meta d’ for “yes” and “no” responses were similar on a ‘pseudo’ detection task, in which the target-absent condition contained not just the absence of a certain physical feature, but the feature was replaced by another feature (unfilled circles). Above we suggested that ‘no’ judgments are somewhat more convoluted or difficult than ‘yes’ judgments, because in the absence of evidence it may be difficult to assess certainty. If this is true, this may only apply to our (true detection) YN memory task and not to our YN visual task, which is akin to ‘pseudo’ detection. This is because unlike with true detection tasks, in our YN visual task subjects didn’t discriminate between the presence or absence of a stimulus; instead they judged whether a stimulus was big or small. Overall, the pattern of results support the hypothesis that it is metacognition after a ‘no’ responses in ‘true’ detection that is causing the problem in limiting our ability to reveal metacognition across task domains.

Our results in large part agree with other similar studies, and may inform their interpretations. As in our Experiment 2 (and Baird et al., 2013), both Baird et al. (2015) and Sadeghi et al. (2017) used YN judgments for their memory task (which were true detection tasks) and 2AFC judgments for their visual task and and found no correlation between metacognitive sensitivity for visual perception and memory. The authors interpret these results as reflecting a genuine lack of correlation or a genuine difference between the task domains, but perhaps the real culprit is the use of a YN task in the memory domain. Likewise, Fitzgerald et al. (2017) also used YN judgments for both their memory task (a true detection task) and visual task (a pseudo detection task) and found no correlation.

Several comparable studies also agree with our Experiment 1 and McCurdy et al. (2013). Samaha and Postle (2017) used multi-choice orientation estimation tasks, which are not 2AFCs but are certainly not genuine YN detection tasks also, and they found a positive correlation between visual and memory metacognitive performance. Faivre et al. (2017) used 2AFCs and found positive correlations for metacognitive efficiency across the visual, auditory, and tactile perceptual domains.

Despite the above agreement, one relevant finding conflicts with our results. As in our Experiment 1 and McCurdy et al. (2013), Morales et al. (2017) required 2AFC judgments for both their visual and memory task. However, they found no correlation across domains for metacognitive efficiency. A likely explanation is that Morales et al. (2017), which included 24 subjects, was underpowered to detect an effect in comparison with our Experiment 1, which included 100 subjects. In support of this possibility, power analyses revealed that given the effect size found in our Experiment 1 with 100 subjects (before subject exclusion; effect size = 0.2470), with alpha set at < .05, power was only .2228 for n=24. This means that even if the effect was actually there, it was unlikely to have been detected given the sample size, so a null finding is unsurprising.

Also in somewhat of disagreement with our results here, Valk et al. (2016) used 2AFC for visual perception and multiple choice questions with three response options for higher-order cognition and found no correlation across these domains. In another study, Garfinkel et al. (2016) used a variety of YN and two-choice tasks (e.g.heart-rate synchronicity detection, tactile grating orientation discrimination, inspiratory resistance detection; none of these were 2AFC tasks) and yet found a positive correlation (unlike in our Experiment 3) between cardiac and respiratory metacognition, but not between either of these domains and tactile metacognition. It is arguable that none of the tasks in Garfinkel et al (2016) are ‘true’ detection tasks, but given the results of Valk et al (2016) it may also be the case that our proposed view regarding YN and 2AFC tasks here only applies to studies of metacognition for certain specific perceptual and memory domains. Alternatively, it could be related to the above mentioned issue of sample size and power. But ultimately, it also leaves open the possibility that our proposed view is incomplete, as the present study may not be definitive - we also need to consider some shortcomings of the present study.

One limitation of our study is that subjects were recruited and completed the tasks online; studies of this nature are relatively new, and therefore one may wonder whether our results are trustworthy. To address this issue, not only did we use catch trials (see Methods), but we took care to weed out subjects with unrealistic results; any participants with d’ lower than .5 or meta d’ less than 0 or percent correct values less than or equal to 60% or greater than or equal to 90% for either or the two tasks was excluded from analyses. Importantly, that the average M Ratio and response times were similar in McCurdy et al. (2013) and our Experiment 1 further suggests that the results of our online experiments are likely legitimate. For McCurdy et al. (2013), mean M Ratio for the visual and memory tasks were 858 ms and 965 ms, respectively, and mean response time was 855 ms and 1,551 ms, respectively; similarly, for our Experiment 1, mean M Ratio for the visual and memory tasks were 821 ms and 1,092 ms, respectively, and mean response time was 424 ms and 1,634 ms, respectively.

Another issue is that our first two experiments were not direct replications of McCurdy et al (2013) and Baird et al (2013), and arguably these two are the studies in question here. One notable difference was that circle stimuli were used for both tasks in the present study, whereas gratings and words were used for the visual and memory tasks, respectively, in the previous studies. While in one sense this might be perceived as a limitation, matching the stimulus type across the tasks was meant to correct a potential confound in the previous studies (as already described), and it is therefore our opinion that it should be viewed as a strength rather than a limitation.

To conclude, historically 2AFC and YN judgments were well-defined task procedures that were clearly distinguished from one another (Macmillan and Creelman, 2004). However, in recent years it has become increasingly common for researchers to label any two choice (discrimination) tasks as 2AFC (Peters et al., 2016). While this may seem like just a terminological issue, the results of our experiments show that conflating 2AFC and YN judgements can lead to substantive consequences. Future studies should avoid this subtle, yet important confound. Because meta-d’ measures are also best suited for 2AFC tasks (Maniscalco & Lau, 2014), we suggest using genuine 2AFC tasks whenever possible. In most instances this is easy to implement: for any two-choice task, we can modify it to present both stimuli in the same trial in temporal succession, or spatially one next to another, and ask the subject to identify the temporal or spatial arrangement of the pair, instead of to tell the identity of a single stimulus. Because one of the two stimuli can be ‘blank’, a two-choice spatial localization task also counts as 2AFC (Macmillan and Creelman, 2004). Therefore, fortunately, the issues discussed here are easy to empirically address in future experiments.

## Acknowledgments

This work was supported by US National Institute of Neurological Disorders and Stroke (NIH R01 NS088628 to H.L.)

